# Storage protein biosynthesis is affected by ionome composition in soybean (*Glycine max* (L.) Merrill) seeds

**DOI:** 10.1101/2025.02.28.640933

**Authors:** Gabriel Sgarbiero Montanha, Lucas Coan Perez, Andrea Lepri, Chiara Longo, Davide Marzi, Eduardo Santos, Felipe Sousa Franco, Gustavo Paparotto Lopes, Nicolas Gustavo da Cruz da Silva, João Paulo Rodrigues Marques, Eduardo de Almeida, Alex Virgilio, Carlos Alberto Pérez, Paola Vittorioso, Sabrina Sabatini, Francisco Scaglia Linhares, Hudson Wallace Pereira de Carvalho

## Abstract

Soybean seeds are a significant source of protein for human and animal nutrition, primarily due to seed storage proteins (SSPs) from the albumin and globulin families, which are predominantly located in protein storage vacuoles within cotyledon cells. This study characterised the dynamics of protein and mineral nutrient accumulation in four soybean genotypes with contrasting protein content—two transgenic (*tg1* and *tg2*) and two conventional (*ct1* and *ct2*)—from the beginning of seed filling (R_5.5_) through to maturity (R_8_) under field conditions. Profiles of globulin SSPs (glycinin and β-conglycinin), as well as the protein and elemental distribution in mature seed cotyledons were examined. Results revealed that genotypes with higher protein content showed increased S and Zn concentrations and a higher glycinin:β-conglycinin ratio. Subcellular analyses further indicated co-localisation of proteins and Zn within cotyledon cells. Our findings reveal a complex association between S and Zn accumulation and SSPs’ biosynthesis, indicating that their availability can limit the SSP content.

**HIGHLIGHT:** Soybean seed genotypes containing higher sulphur (S) and zinc (Zn) content in the cotyledonary cells exhibit a distinct storage proteins profile by increasing the abundance of sulphur-amino acids rich globulins.

## 1. INTRODUCTION

Soybean (*Glycine max* (L.) Merrill) is a crucial source of protein for human and animal nutrition. With seeds accumulating *ca*. 40 wt.% protein (Dashiell, 2005), soybeans contribute to more than 70% of the world’s protein meal output, significantly surpassing other species such as rapeseed (*Brassica napus*) and sunflower (*Helianthus annuus*) (American Soybean Association, 2024). Additionally, soybean seeds are an essential source of vegetable oil, used in food, various industrial applications, and biofuel production (Ağbulut *et al*., 2024).

Seed storage proteins (SSPs) constitute up to 80 wt.% of the soybean seed proteome and, consequently, form the bulk of the protein meal (Murphy, 2008; Song *et al*., 2016). SSPs are synthesised in the endoplasmic reticulum (ER) and transported *via* a tightly regulated Golgi-mediated process to the protein storage vacuoles (PSVs), a specialised group of vesicles predominantly found in the seed’s cotyledon cells (Li, 1993; Herman and Larkins, 1999). They serve as a source of amino acids for germinating seeds and seedlings (Derbyshire *et al*., 1976; Aarabi *et al*., 2021). In soybeans, SSPs are classified according to their solubility into water-soluble albumins and salt-soluble globulins (Day, 2013).

Globulins comprise *ca*. two-thirds of the total SSPs and consist of two main proteins: glycinin (legumin), a hexamer of acidic-basic polypeptide chains linked by disulfide bonds, and β-conglycinin (vicilin), a trimer composed of glycosylated α, α’, and β peptides (Nielsen, 1985; Harada *et al*., 1989; Nielsen *et al*., 1989). In contrast, albumins are present in lower concentrations and are primarily low molecular weight proteins smaller than 20 kDa, while globulins range from 25 to 72 kDa (Murphy, 2008).

From a nutritional standpoint, despite its balanced composition, soybean seeds are notably deficient in sulphur amino acids (SAAs) such as methionine (Met) and cysteine (Cys), either essential or conditionally essential for monogastric animals, including humans (Wang *et al*., 2022). Cys and Met typically constitute only 1.4–1.6% of soybean meal amino acid profiles (Krishnan and Jez, 2018).

The SAA scarcity is attributed to the higher abundance of the β-conglycinin β-peptide, which lacks methionine and contains one or no cysteine residues, compared to glycinin, which has 2–5 methionine and 2–6 cysteine residues per subunit (Imsande, 2001; Zarkadas *et al*., 2007). Consequently, exogenous supplementation of these amino acids is frequently required in soybean meal-based feeds (Krishnan, 2015; Krishnan and Jez, 2018; Rushovich and Weil, 2021). Projections from the United Soybean Board’s Better Bean Initiative (BBI) indicate that a 50% increase in SAA is necessary to meet the optimal requirements of the poultry feed industry (Krishnan and Jez, 2018).

However, as with many other agronomic traits in soybeans, such as yield, seed size, and oil content, protein content, and composition are controlled by quantitative trait loci (QTLs), which are regulated by multiple genes. Several QTLs have been identified in close genomic regions, leading to potential linkage, pleiotropy, and environmental influences (Qi *et al*., 2011; Borja Reis *et al*., 2021).

Notably, while breeding programmes have achieved an impressive 21% per decade increase in soybean yield over the last 50 years, this has been accompanied by a 1.1% per decade decrease in protein content (de Borja Reis *et al*., 2020; Umburanas *et al*., 2022; Montanha *et al*., 2024). Similarly, the concentration of amino acids followed a similar trend in soybean genotypes released between 1980 and 2014 in the United States (de Borja Reis *et al*., 2020), suggesting that the decline in total protein content did not significantly affect the balance of SSPs in soybean seeds. In other words, the decrease in protein does not seem to be selective towards a specific type of protein, but it affects the proteome as a whole.

Soybean SSPs are synthesised primarily from the seed-filling phase (R6 stage) onwards (Herman and Larkins, 1999; Montanha *et al*., 2023). Despite the complexity of SSPs profile regulation, one should keep in mind that its accumulation is also intrinsically associated with nutrient availability. In a previous field trial from our group, SSP accumulation peaked during the mid-R6 stage, coinciding with increased sulphur (S) and magnesium (Mg) concentrations in seeds (Montanha *et al*., 2023). A screening of 95 soybean varieties from Brazil revealed a complex association between total protein and S, zinc (Zn), and manganese (Mn) content (Montanha *et al*., 2024).

Sulphur amino acids account for around 80% of all organic S in plants (Borja Reis *et al*., 2021). Besides its structural role in Met and Cys, S is responsible for forming disulfide bonds between the acidic and basic polypeptides of glycinin. Interestingly, increased β-conglycinin β-peptide levels, which lack SAAs, have been observed in *Petunia hybrida* seeds grown under S deficiency (Fujiwara *et al*., 1992), further highlighting the potential role of S in SSP synthesis. Additionally, Mg and Mn are involved in maintaining PSVs in seeds (Monma *et al*., 1992; Ferreira *et al*., 1999; Santos *et al*., 2012; Madsen and Brinch-Pedersen, 2020), while Zn is essential for protein biosynthesis (CAKMAK *et al*., 1989) and acts as a cofactor for methionine synthase in plants (Eckermann *et al*., 2000).

These data suggest nutrient availability may limit SSP biosynthesis in soybean seeds. Therefore, ensuring an adequate nutrient supply to soybean plants could influence SSPs’ concentration and composition. However, the precise role of these nutrients in SSP biosynthesis remains poorly understood. Therefore, the present study aims to characterise the dynamics of protein and mineral nutrient accumulation in soybean genotypes grown under field conditions.

## 2. MATERIALS AND METHODS

### Cultivation

A field experiment using two transgenics, *i.e.*, RSF ‘Lança’ (58I60RSFIPRO, BrasMax, Brazil) and RSF ‘Zeus’ (55157RSFIPRO, BrasMax, Brazil), and two conventional, *i.e.*, BRS232 and BRS282, both provided by Embrapa Soja (Brazil), hereby named as *tg1*, *tg2*, *cv1*, *cv2*, soybean varieties with contrasting total protein contents, were carried out between November 16^th^ 2021 and March 15^th^ 2022 in an agronomic research station at the municipality of Iracemápolis, São Paulo State, Brazil (22°38’S, 47°30’W, 564 m elevation, **Fig. S1-A**). Prior the experiment onset, the total protein content in the seeds was determined through the Dumas method, and the average obtained values were 39, 41, 40, 38% wt. for the *tg1*, *tg2*, *cv1*, *cv2*, respectively. Shortly before the sowing, the seeds were coated with 25 g L^-1^ pyraclostrobin, and 250 g L^-1^ fipronil (StandakTop, BASF, Brazil) at 2 mL kg seed^-1^, 30 g L^-1^ K_2_O, 6 g L^-1^ Co, 120 g L^-1^ Mo, 12 g L^-1^ Ni (Up! Seeds, ICL, Brazil) at 3 mL kg seeds^-1^, graphite powder at 5 g kg seeds^-1^, and inoculated with a 5x10^9^ mL^-1^ minimum colony forming units (CFU) *Bradyrhizobium elkanu* and *Bradyrhizobium japonicum* (SEMIA 5019 and 5079 strains, respectively) solution (MasterfixL, Stoller, Brazil) at 2 mL kg seeds^-1^. The 2.8 ha experimental plot encompassed four *ca*. 350 m x 10 m lines, in which the seeds were sown at a 3-cm depth using a planter with six lines with a 20-cm distance from each other and 16 seed m^-1^ density. A composite soil sample collected from 0 to 0.2 m depth before the experiment exhibited the following chemical characteristics: pH: 5.57; organic matter: 20.28 g dm^−3^; base saturation: 77.03%; S: 29.36 mg dm^−3^; P: 44.69 mg dm^−3^; K: 4.73 mmol_c_ dm^−3^; Ca: 56.32 mmol_c_ dm^−3^; Mg: 12.14 mmol_c_ dm^−3^; Al: 0 mmol_c_ dm^−3^; H+Al: 21.68 mmol_c_ dm^−3^; Cu: 1.85 mg dm^−3^; Fe: 12.08 mg dm^−3^; Zn: 3.07mg dm^−3^; Mn: 3.90 mg dm^−3^; B: 0.46 mg dm^−3^. The average temperature and accumulated rainfall during the experiment were 25°C and 541.2 mm, respectively.

### Assessment of soybean seed development

Soybean pods from R_5.5_, *i.e.*, the beginning of the formation of the seeds, to R_8_, *i.e.*, mature seeds phenological stages, were manually collected in the early hours of the morning at the median height of soybean plants within 6 blocks within *ca*. 1 m^2^ areas randomly selected within each experimental plot until reaching suitable mass, and the sampled regions were tagged using a digital GPS and not used in the further collections (**Fig. S1-BC**). At the mature R_8_ stage, all the soybean plants within 6 blocks of 18 m^2^ each were harvested using a brush-cutter (**Fig. S1-D**), and the soybean seeds were obtained through a mobile thresher (**Fig. S1-E**).

The freshly collected soybean pods were weighed and then face-scalped to assess seed structure and development using a high-resolution scanner (HP model 2136, HP, Brazil). The seed length was determined using Fiji open-source software (version 2.1.0/1.53c), then they were dried in a fan-circulated oven (Tecnal, TE-394/2-MP, Brazil) at 60 °C for at least 72-h or until reaching constant mass for determining the H_2_O content and the 1000-seed dry mass or kept in a dry chamber at 10°C for remaining analyses.

For the mature seeds, these values were also used for determining the yield index, calculated according to Equation 1:

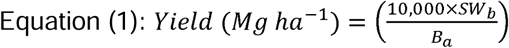

Where: SW_b_ is the total seed dry mass collected from a certain block, in Mg, and B_a_ is the total area of the block, *i.e.*, 18 m^2^.

### Assessment of mineral and total protein composition through seed development

The seeds were manually removed from the pods, then also weighed and either dried in a fan-circulated oven (Tecnal, TE-394/2-MP, Brazil) at 60°C for at least 72-h or until reaching constant mass for determining the mineral composition or kept in the low-humidity chamber at 10°C for the remaining analyses. The oven-dried soybean seeds were ground using a blade miller (Cadence, model MDR302, Brazil) for two cycles of 30 s each. The quantification of protein in soybean seeds was carried using the Dumas official methodology from the Association of Official Analytical Chemists (AOAC, 1997), according to the procedure described by Montanha *et al*., 2024. For determining the mineral composition, 100 mg of the ground seeds were digested in a TFM vessels through a microwave-assisted system (Ultrawave, Milestone, Sorisole, Italy) sample masses of 100 mg were accurately weighed directly in the TFM vessels, and 6.0 mL of 20% v v-1 HNO3 (Merck, USA) and 1.0 mL of 30% w w-1 of H2O2 (Dinâmica, Brazil) were added. The 4-step heating program was: i) 5 min ramp to 160 °C, ii) 2 min hold at 160 °C, iii) 5 min ramp to 230 °C and iv) 15 min hold at 230 °C. After reaching room temperature, the digested solutions were transferred to 50 mL Falcon tubes and the final volumes were made up to 25 mL with ultrapure water. The resulting solutions were analysed by inductively coupled plasma optical emission spectrometry (ICP OES), using an iCap 7400 Duo-spectrometer (Thermo Scientific, Waltham, MA, USA). The instrument’s operating conditions were: RF Power at 1200 W, plasma gas flow at 12 L min^-1^, nebulizer gas flow at 0.6 L min^-1^, auxiliary gas flow at 0.5 L min^-1^, 15 s of integration time and 60 rpm of pump rotation. The analytes and selected wavelengths were: Ca (422.673 nm), K (766.490 nm), Mg (285.213 nm), B (249.773 nm), P (213.618 nm), S (180.731 nm), Cu (324.754 nm), Fe (238.024 nm), Mn (257.610 nm) and Zn (213.856 nm). The accuracy of the digestion procedures and the ICP OES analysis was determined by measuring the concentration of the analytes certificate reference materials. The recoveries ranged from 85 – 118%. All measurements were accomplished using 4 independent replicates.

### Assessment of protein storage vesicles (PSVs) through light microscopy and synchrotron-based computed tomography (µ-CT)

For the light microscopy analyses, the cotyledonary tissues of mature soybean seeds from the four soybean varieties were manually excised using a scalpel. The samples were fixated, dehydrated, embedded in historesin (Leica, Germany) and cut using a rotatory microtome (Leica, model RM2125 RTS, Germany), according to the methodology described byMarques & Soares, 2022. The resulting 10 µm-thick sections were stained for 30 s with a 1% v v^-1^ xylidine ponceau solution, then washed in 3% v v^-1^ acetic acid for 15 min, mounted in a glass slide and directly observed in a microscope (Zeiss, Axion Observer, Germany) under a 20x magnification. In each genotype, the measurements were accomplished using 10 independent replicates. The PSVs number and area were quantified using the Fiji (version 2.1.0/1.53c) software.

The µ-CT analyses were carried out at the Biomedical Imaging and Therapy (BMIT-BM) beamline of the Canadian Light Source, in Saskatoon, Canada. In this regard, the median region of mature soybean seed’s cotyledonary were cross-sectioned and exposed to a 20 horizontal x 4 mm-vertical X-ray beam operating at 30 keV with a 800 µm Al and 100 µm Ag primary filters selected. The imaging field was 240 x 240 µm and 1200 projections were acquired over a 360° rotation of the sample with a 0.3 µm pixel size and a 650 ms per projection exposure time. The three-dimensional images were reconstructed and the PSVs were isolated for the volume analysis using the Avizo Software (ThermoFischer Scientific, USA), as shown in **Fig. S2**. At least 3 independent biological replicates of each genotype were employed.

### Assessment of globulins profile through SDS-Page

Glycinin and β-conglycinin were extracted from soybean seeds according to the procedures described by Wei et al., 2020, with minor modifications. Briefly, mature seeds from 58I60 RSF IPRO, 58I60 RSF IPRO, BRS232, and BRS284 varieties (hereby referred to as *tg1*, *tg2*, *ct1*, and *ct2*, respectively) were powdered by manually grounding to a fine powder using a mortar containing liquid nitrogen. Thereafter, 10 mg of each sample was transferred to a centrifuge vial, followed by the addition of 1 mL of SDS-sample buffer solution (60mM tris-HCl at pH 6.8, 2% sodium dodecyl sulphate, 10% glycerol, 2.5% β-mercaptoethanol), vortexing for 10 min at room temperature, and boiling for 5 min. The samples were centrifuged at 15,000 g for 5 min, and the supernatant was collected in a fresh vial. Seed proteins were resolved using a one-dimensional SDS-PAGE electrophoresis kit (Bio-Rad, USA). Precisely 5 µL of each sample was added to 2 µL of Laemmly 1X tracking die (Bio-Rad, USA) and loaded onto a 12% acrylamide gel (TGX Stain-Free™ FastCast™ Acrylamide Kit, Bio-Rad, USA) covered with running buffer solution (3.03% tris base, 14.4% glycine, 1% sodium dodecyl sulfate). The run was carried out at a constant of 120 V/gel until the tracking dye reached the bottom of the gel. After that, the gels were carefully removed from the cassette and stained with 0.25% Coomassie Blue solution (0.25% Coomassie Brilliant Blue R250, 90% methanol: H_2_O (1:1 v/v), 10% glacial acetic acid) for 30-min, then destained overnight (60% H_2_O, 30% methanol, 10% acetic acid), and imaged at a ChemiDoc MP Imaging System (Bio-Rad, USA). The experiments were accomplished using three technical replicates and three independent biological replicates of each seed variety. The recorded values of each selected protein bands were subtracted from their respective background values and normalised by the total protein bands of each lane using the ImageLab (Version 6.1.0, Bio-Rad, USA) software.

### Combined FTIR and synchrotron-based XRF mapping of soybean seed cotyledon

Mature soybean seed samples from the four varieties were manually peeled and trimmed using a razor blade, then directly mounted on an ultramicrotome (Leica, model UC6, Germany). The median region of the cotyledons (*i.e.*, those that accumulate most storage proteins) was cross-sectioned using a glass knife. The resulting 10-μm-thick cross-sections were immediately transferred to a Si/Au wafer and used for sequential Fourier transform infrared spectroscopy (FTIR) and X-ray fluorescence spectroscopy (XRF) mappings, as detailed in **Figure S3**. The sample integrity was assessed by direct observation of the samples through transmitted light microscopy (Nikon, model Eclipse LV100ND, Japan), or they were transferred to a glass slide, stained with toluidine blue (TBO, Sigma-Aldrich, Germany), and observed through light microscopy (Zeiss, Axion Observer, Germany).

The Fourier transform infrared spectroscopy mappings were carried out at the benchtop station (Agilent Cary 600 Series, USA) of the Imbuia beamline at the Brazilian Synchrotron Light Laboratory, Campinas, Brazil. The maps were recorded through 128 sequential scans in absorbance mode within 420 × 420 µm^2^ areas with a lateral resolution of 5.5 × 5.5 µm. The wavenumber ranged from 900 to 3000 cmL¹, with a 4 cmL¹ resolution. The intracellular quantification of proteins, lipids, and pectins was carried out by integrating the 1586–1710 cmL¹, 1714–1792 cmL¹, and 914–1141 cmL¹ regions, respectively, with at least 20 regions selected in each sample, as detailed in **Figure S4**.

The X-ray fluorescence spectroscopy mappings were carried out at the Tarumã endstation of the Carnauba coherent X-ray nanoprobe beamline at the Brazilian Synchrotron Light Laboratory, Campinas, Brazil. The Si/Au wafer containing the samples were placed into a metallic XRF sample holder, covered with carbon-fibre tape, and transferred to the beamline. This beamline features a horizontal deflection four-bounce crystal monochromator (4CM), two four-element silicon drift detectors (Vortex-ME4, Hitachi High-Technologies Science America, USA), and a Kirkpatrick−Baez (KB) achromatic optic providing a 150 nm-wide X-ray focused beam spot. The XRF maps were recorded under room-temperature condition in flyscan mode at an energy slightly higher than the Zn K-line excitation energy (10,300 eV), through 400 × 400 µm panoramic images. The data were processed using PyMCA software (version 5.6.3, ESRF Software Group, France). The elemental intensities detected in each pixel unit were normalised by the ring current (IL) and fitted according to the beamline instrumental parameters. The experiment was carried out using at least three independent biological replicates. As shown in **Figure S5**, the measurements using the Si/Au substrate enabled the detection of K, Ca, and Zn.

The Zn intensities within the cotyledon cells were quantified in at least 20 cells per sample using Fiji software (version 2.1.0/1.53c). Only the fitted elemental intensities above the fitting background, i.e., those > 0, were considered valid. Additionally, the Ca (used as a marker of the cell wall) and Zn intensities were quantified separately in the cell wall and cytoplasm regions of three randomly selected samples to certify that the obtained intensities were not instrumental artefacts, as shown in **Figure S6**.

### Statistical analyses

The normality of the data was assessed through the Shapiro-Wilk test at 0.05 level. The differences between each group were evaluated either through *t*-test or Kruskal–Wallis one-way analysis of variance followed by Dunn’s multiple comparisons test at a 95% confidence level (*p* < 0.05). Two-variable correlations were determined through Spearman’s rank correlation coefficient test at a 95% confidence interval, followed by a K-means hierarchical clustering. All analyses were carried out using Prism (version 9.4.0, USA) and RStudio (version 1.4.1106 "Tiger Daylily", USA; packages ‘*stats*’ and ‘*ComplexHeatmap*’) software.

## 3. RESULTS AND DISCUSSION

To assess the dynamics of mineral nutrients and protein accumulation in soybean seeds, we evaluated four soybean genotypes cultivated under field conditions: two conventional and two transgenics, *i.e.*, bearing resistance to glyphosate herbicide and caterpillars (Silva *et al*., 2019). These genotypes are commercially employed in Brazil, and exhibit contrasting total seed protein content, as previously described (Pereira, 2024; Umburanas *et al*., 2022).

**Figure 1** presents the characterisation of soybean seed development from the beginning of seed filling (R_5.5_) to its full maturity (R_8_) (Fehr and Caviness, 1977), revealing that both seed length and fresh mass increased exponentially until reaching the highest values at R_6_ developmental stages, then started decreasing together with a sharp reduction in H_2_O content. In contrast, the seed dry-mass values reached the maximum value at the maturity stage (R_8_). All genotypes exhibited a very similar trend for all variables assessed. Although it indicated an earlier onset of the seed dehydration phase in both transgenic genotypes (*tg1*, *tg2*) than in the conventional ones (*ct1* and *ct2*), mature seeds showed virtually the same H_2_O content and both fresh and dried mass.

**Figure 1.**
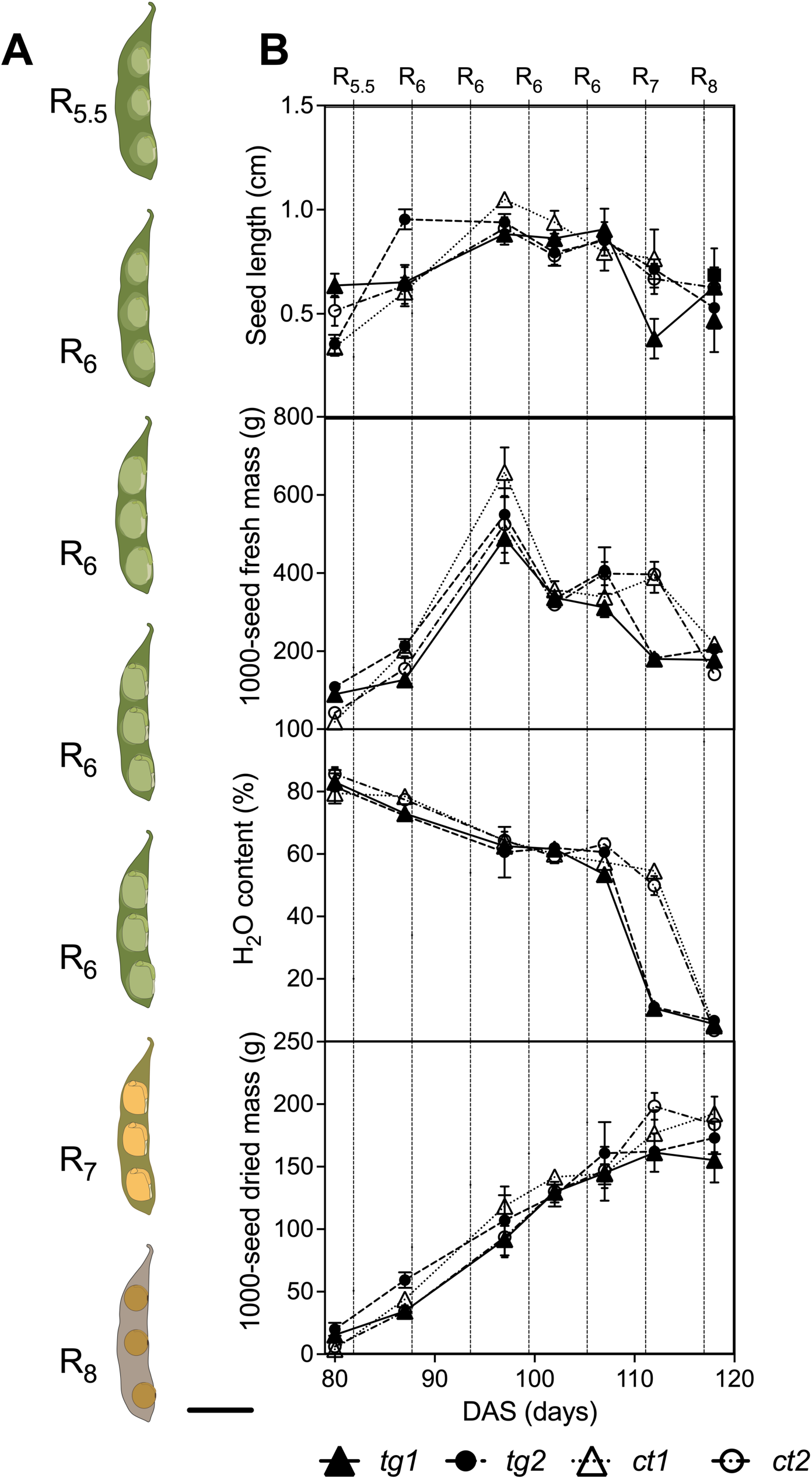
Characterisation of soybean seeds development. Schematic representation of cross-sectioned soybean pods and seeds (a), seed length, 1000-seed fresh mass, H_2_O content, and 1000-seed dry mass (d) recorded in four soybean varieties, *tg1*,*tg2*, *ct1*, and *ct2*, collected in field conditions from the beginning of the seed filling (R_5.5_) up to maturity (R_8_) stages. All quantitative values represent the mean ± standard error (SE) recorded using at least 3 independent replicates with 20 seeds each. Scale: 1 cm.

After fertilisation, the zygote and maternal cells undergo a complex differentiation process into seed coat, embryo, and cotyledon tissues spanning between the R_3_ and R_5.5_ developmental stages (Lin *et al*., 2017). During the R_6_ stages, storage compounds, such as proteins, oil, and carbohydrates, accumulate in these tissues before the desiccation (R_7_) and quiescence/mature (R_8_) stages (Montanha *et al*., 2023). This dynamic highlights the biometrical data obtained, *i.e.* both seed length and fresh mass maximum values were recorded during the R_6_, and the maximum 1000-seed dry mass is found at the mature R_8_ stage, where the H_2_O reaches the lowest values (**Fig. 1-B**).

Furthermore, **Fig. 2** shows the concentration of total protein, as well as macro (K, P, Ca, S, and Mg) and micronutrients (Fe, Zn, Mn, B, and Cu) in soybean seeds throughout their development. Regardless of the genotype, seeds exhibited a very low protein content, that is ca. 23 wt.% at the beginning of the seed-filling stage (R_5.5_), which consistently increased during the R6 stages and reached its highest values, *i.e.*, 35-39 wt.% at maturity (R_8_). A similar trend was observed for the S and Mg concentrations, whereas a significant reduction was recorded for all other nutrients as a function of the seed developmental stage. The accumulated values as a function of the seed yield are presented in **Figures S7-S8**.

**Figure 2.**
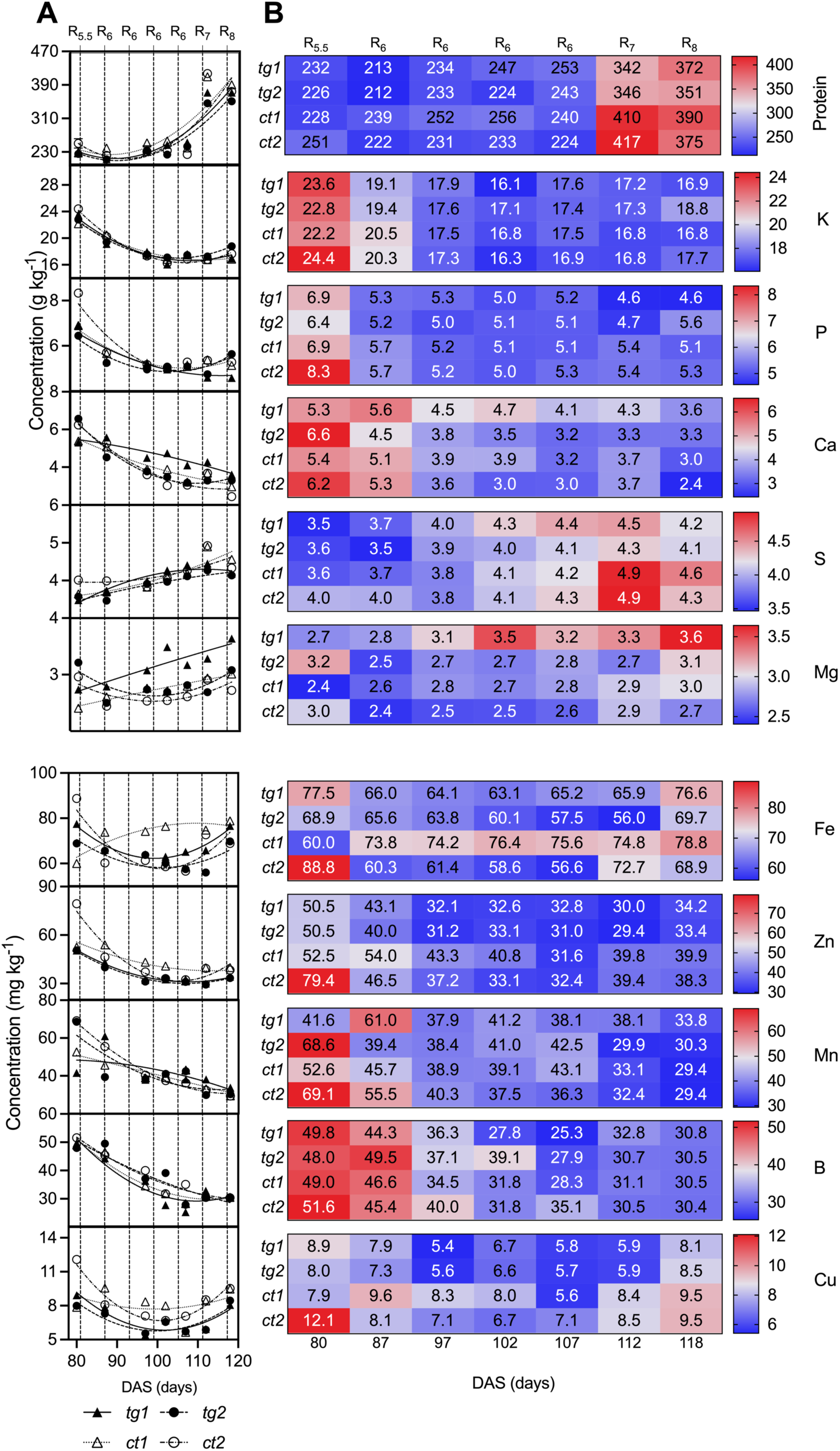
Characterisation of soybean seeds development. Schematic representation of cross-sectioned soybean pods and seeds (a), seed length, 1000-seed fresh mass, H_2_O content, and 1000-seed dry mass (d) recorded in four soybean varieties, *tg1*, *tg2*, *ct1*, and *ct2*, collected in field conditions from the beginning of the seed filling (R_5.5_) up to maturity (R_8_) stages. All quantitative values represent the mean ± standard error (SE) recorded using at least 3 independent replicates with 20 seeds each. The data were fitted according to an exponential equation.

SSPs from the albumin and, noteworthy, globulin families compose more than 70% of the soybean seed proteome and play a vital role as a source of amino-form nitrogen, as well as S, to the germinating seeds and early seedlings (Montanha *et al*., 2023). These proteins are synthesised within the cotyledonary cells by the ER and transported *via* the Golgi apparatus to the PSVs, a specialised vesicle most abundantly found within the seed’s cotyledon cells (Li, 1993; Herman and Larkins, 1999).

In addition to molecular and environmental factors, several pieces of evidence suggest that SSPs biosynthesis during soybean seed development also depends on mineral nutrients (Montanha et al., 2023a; Montanha et al., 2024). This is reinforced by Spearman’s rank correlation coefficients between the total protein and mineral nutrients presented in **Fig. 3-A**, which reveals a strong positive association between protein, S, and Mg concentrations in soybean seeds as a function of their development, which is higher for *ct1* and *tg1* than for *ct2* and *tg2* genotypes, respectively, as shown by the k-means hierarchical clustering. Additionally, **Fig. 3-B** presents the ratio between the protein and mineral nutrient concentrations recorded at the onset of the seed filling (R_5.5_) and mature (R_8_) stages and shows that *ct1* and *tg1* also presented a higher relative concentration of protein, S, and Mg throughout soybean seed development, reinforcing the association between protein, S, and Mg shown in **Fig. 3-A**. Similar results were obtained in a previous field trial (Montanha *et al*., 2023), reinforcing the importance of these mineral nutrients in SSP accumulation. In **Fig. 4** the yield and protein content of mature soybean seeds is presented.

**Figure 3.**
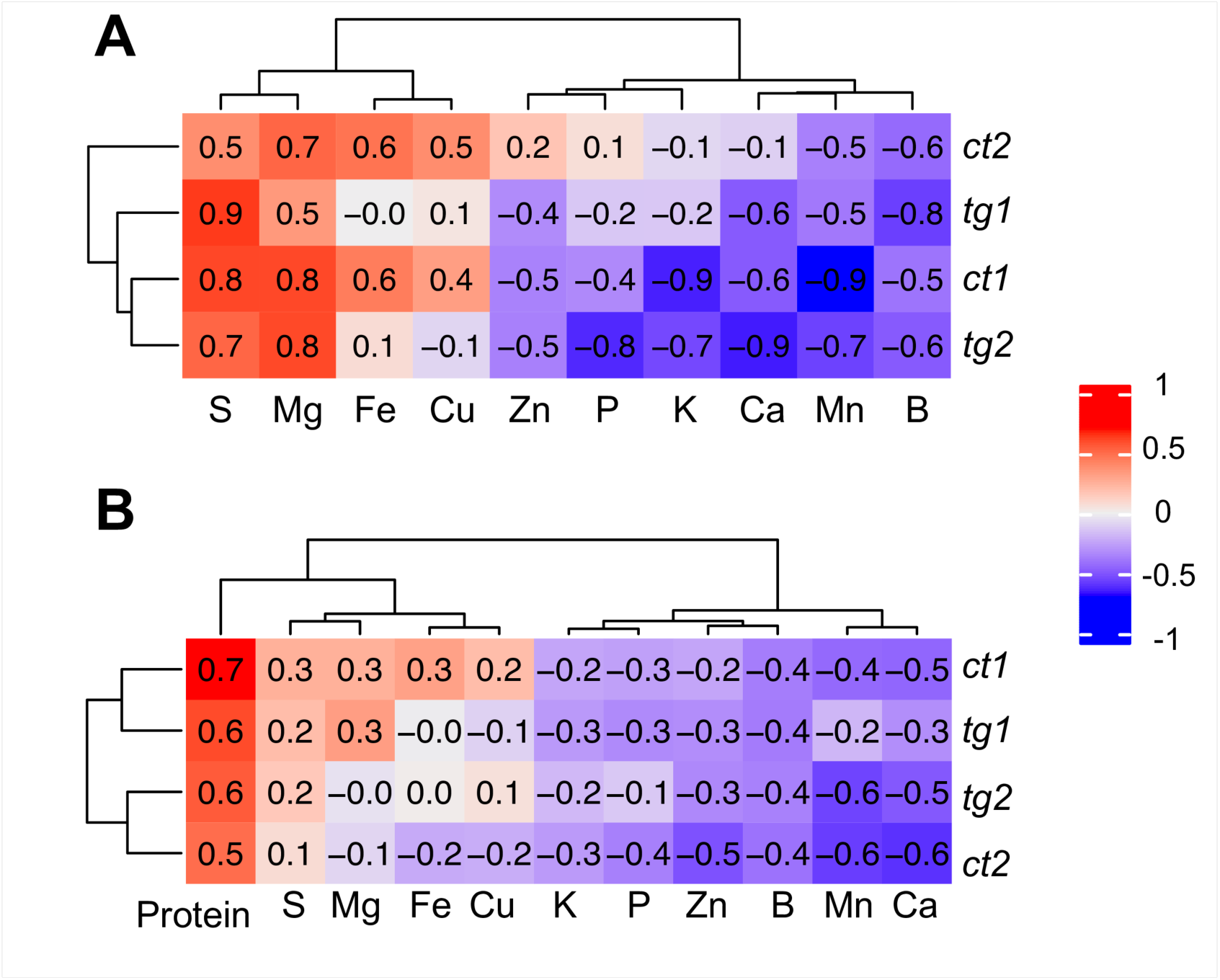
Heatmap of the Spearman’s rank correlation coefficients recorded between the concentration of protein and mineral nutrients in soybean seeds from the *tg1*, *tg2*, *ct1*, and *ct2* genotypes throughout the beginning of the seed filling (R5.5) up to maturity (R8) stages (a), and their ratios at the initial (R5.5) and final (R8) stages observation. The tests were conducted at a 95% confidence interval followed by k-means hierarchical clustering.

**Figure 4.**
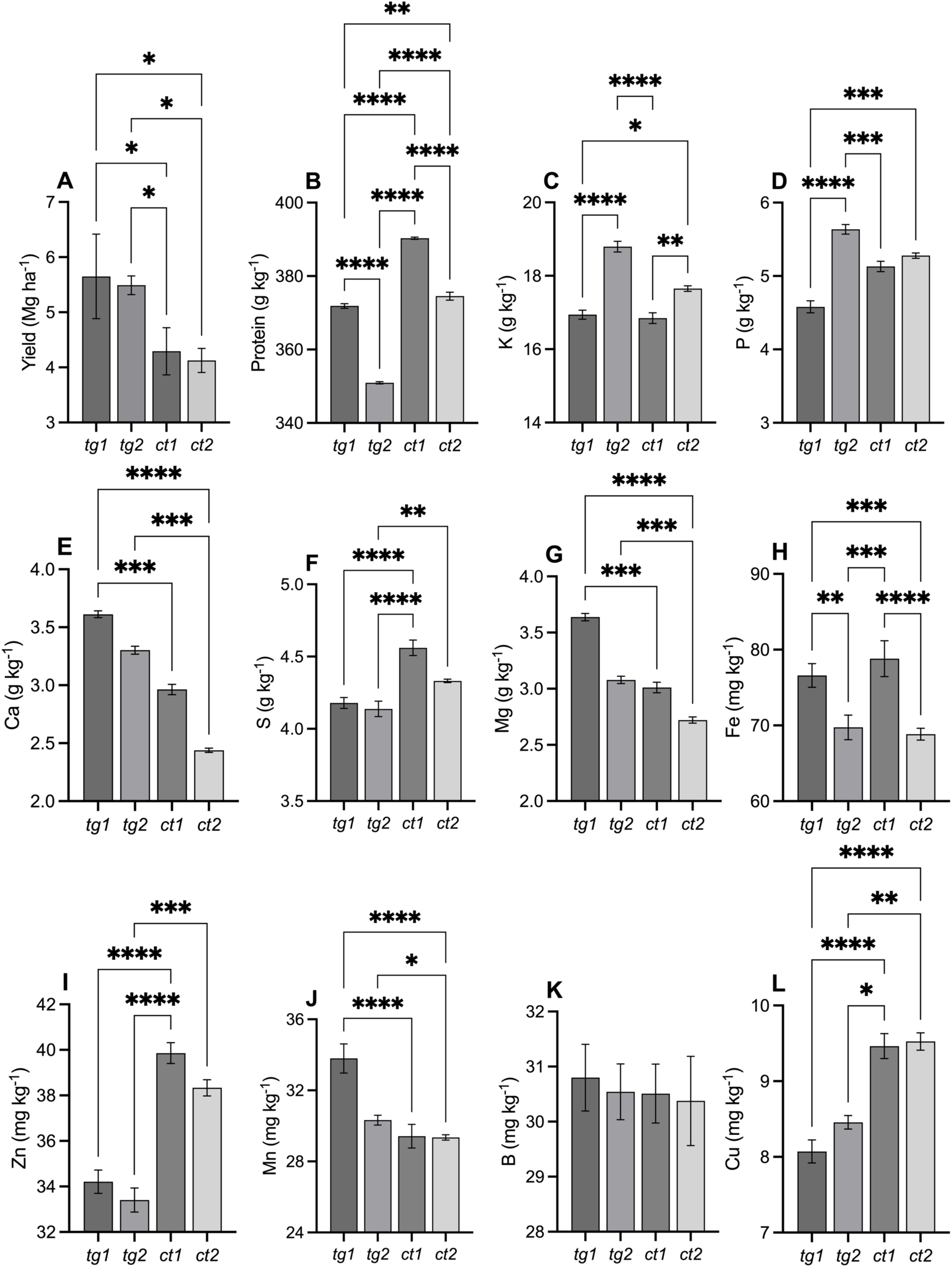
Yield, total protein, K, P, Ca, S, Mg, Fe, Zn, Mn, B, and Cu concentration recorded in mature (R_8_ stage) soybean seed varieties, *tg1*, *tg2*, *ct1*, and *ct2*, collected in field conditions. Data represent the mean ± standard error (SE) recorded using at least 3 independent replicates. Data were subjected to Kruskal–Wallis one-way analysis of variance followed by Dunn’s multiple comparisons test at 0.05 level. *, **, ***, *****= *p* <0.05, *p* <0.01, *p* < 0.001, *p* < 0.0001.

Interestingly, a higher yield was observed for the transgenic genotypes (*tg1* and *tg2*) compared to the conventional genotypes (*ct1* and *ct2*) (**Fig. 4-A**), whereas the opposite was found for the protein content (**Fig. 4-B**). In addition, mature *ct1* and *ct2* seed genotypes exhibited higher S (**Fig. 4-F**), Zn (**Fig. 4-I**), and Cu (**Fig. 4-L**) concentrations than *tg1* and *tg2*. These observations led to the conclusions that the lower the yield, the higher the protein content, S, Zn, and Cu concentrations in soybean seeds. In a previous screening of 95 soybean genotypes collected from several regions in Brazil, we also identified a positive association between the S, Zn, Mn, and total protein concentration (Montanha *et al*., 2024). Taken together, these shreds of evidence corroborate that soybean seeds with contrasting protein concentrations exhibit distinct ionomes.

As SAA accounts for *ca*. 80% of all organic S in plants (Borja Reis *et al*., 2021), and S is also involved in the disulphide bonds of glycinin’s acidic-basic polypeptides (Murphy, 2008), one could hypothesise that the higher the S concentration, the higher the glycinin content in soybean seeds. Thus, globulin proteins were extracted from mature soybean seeds and resolved through one-dimensional SDS-Page electrophoresis, as shown in **Fig. 5-A**. The densitometric analyses of the protein bands revealed that summarised β-conglycinin subunits were slightly higher in the *tg1* and *tg2* genotypes compared to the *ct1* and *ct2* ones, respectively (**Fig. 5-B**). An opposite trend was observed for summarised glycinin subunits (**Fig. 5-C**). As a result, a higher glycinin:β-conglycinin ratio was found for the conventional genotypes (*ct1* and *ct2*) compared to the transgenic ones (*tg1* and *tg2*), as shown in **Fig. 5-D**, thus suggesting a higher abundance of glycinin over β-conglycinin in the seeds exhibiting significantly higher S and Zn concentrations. The independent biological replicates, as well as intensities for the individual glycinin’s acid and basic subunits and β-conglycinin’s α, α’, and β subunits, are presented in **Fig. S9-S10**, respectively.

**Figure 5.**
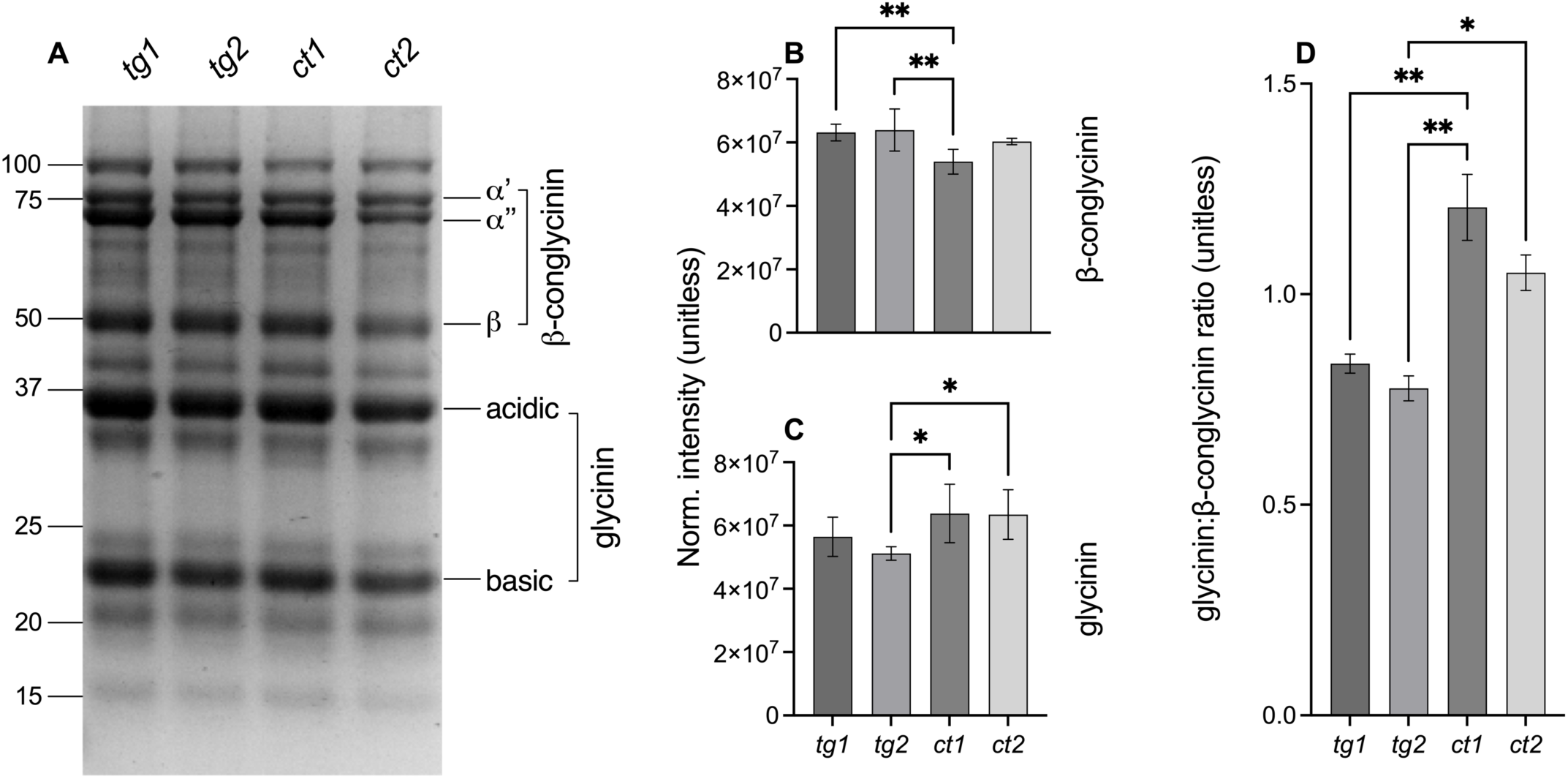
SDS-PAGE of glycinin and β-conglycinin from 4 soybean seed varieties exhibiting contrasting protein contents. Normalised intensities of the β-conglycinin (a) and glycinin (d-e) proteins in the 4 soybean seed varieties. Glycinin: β-conglycinin ratio. Data indicate the mean and the standard error from three biological replicates. The values were compared through one-way ANOVA followed by Tukey’s *post hoc* test at a 95% confidence interval (*p* < 0.05). *, **, ***, *****= *p* <0.05, *p* <0.01, *p* < 0.001, *p* < 0.0001.

Moreover, Zn is associated with more than 300 enzymes and plays a crucial role in key biochemical processes, such as transcription and translation, and hence, protein biosynthesis, as well as antioxidant and catabolic metabolism (CAKMAK *et al*., 1989). Notably, Zn is also a co-factor for the methionine synthase in plants (Eckermann *et al*., 2000), suggesting that the increased Zn content observed in the soybean seed genotypes with high total and glycinin protein regards to its role on SAA biosynthesis. Therefore, we have explored a combined FTIR and synchrotron-based XRF strategy to assess whether the intracellular Zn distribution is associated with protein distribution in soybean cotyledon cells, *i.e.*, where SPVs containing the SSPs are found (Herman and Larkins, 1999).

Figure **6-A** presents the spatial distribution of the amide I, *i.e.* the most intense absorption band in proteins as well as lipids, and pectins FTIR bands (Lee *et al*., 2018; Ji *et al*., 2020; Bhagia *et al*., 2022) obtained in the cross-sectioned soybean seed cotyledon cells. It unveils that most of the pectin intensities is found at the cell wall, whereas the amide I band is mostly found as hotspots within the cells. Conversely, although the distribution of the lipid appears to be more consistent at the cell walls, its intensities were found scattered throughout the intracellular region, which is likely due to the presence of characteristic carbonil (C=O) group vibration (1714–1792 cmL¹) region (Lee *et al*., 2018) at both cellular membrane phospholipids and oil reserve bodies (Li *et al*., 2022).

**Figure 6.**
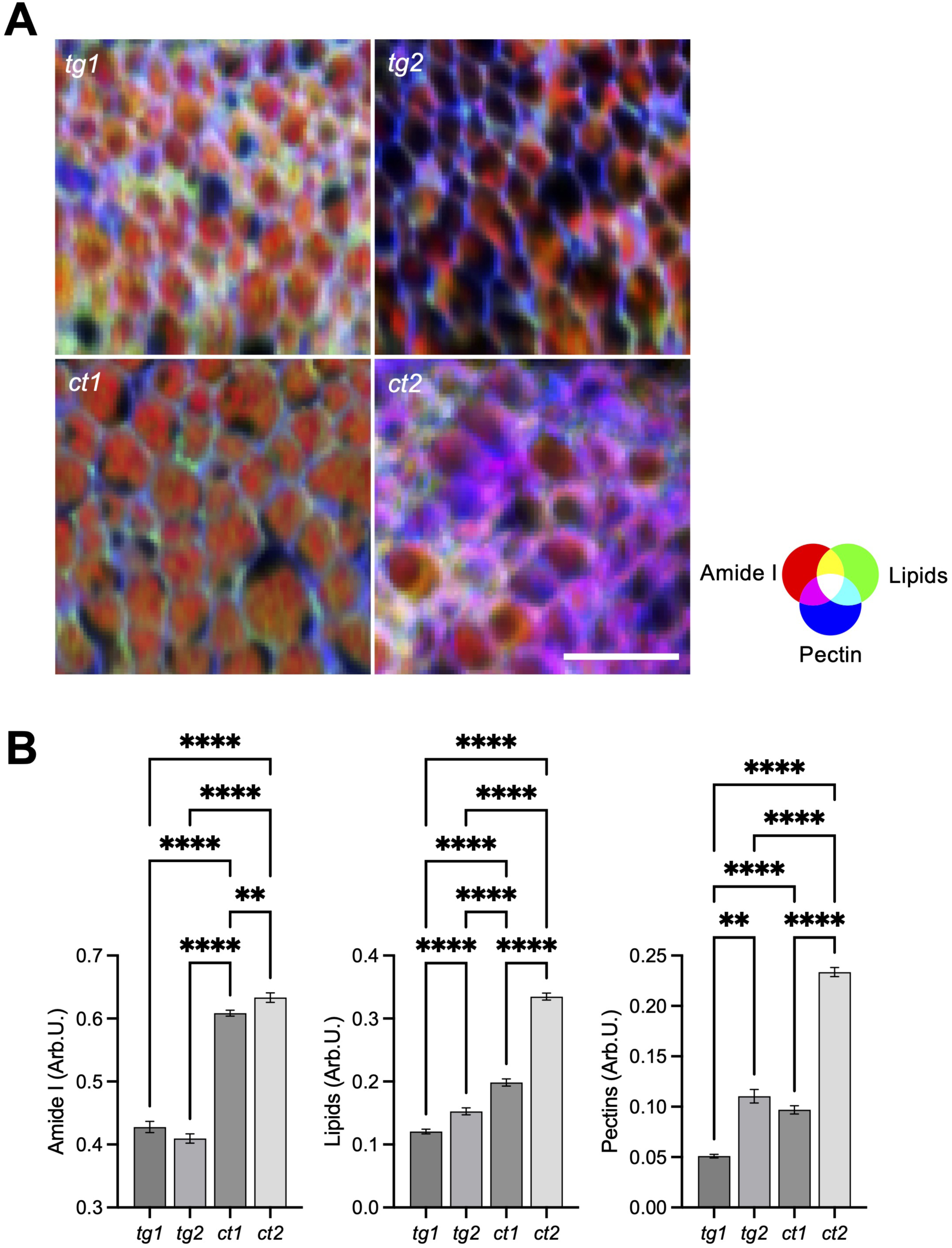
Spatial distribution (a) and average intensities (b) of integrated amide I (red), lipids (green), and pectins (blue) FTIR bands recorded in the cotyledon tissue of mature (R_8_ stage) soybean seed cross-sections from varieties, *tg1*, *tg2*, *ct1*, and *ct2*, collected in field conditions. The bars represent the mean ± standard error (SE) recorded using at least 20 intracellular regions of three independent biological replicates. Data were subjected to Kruskal–Wallis one-way analysis of variance followed by Dunn’s multiple comparisons test at 0.05 level. *, ** = *p* <0.05, *p* <0.01. Scale:100 *μ*m

The subcellular quantification shows that the amide I intensities were significantly higher in the cells from the *ct1* and *ct2* genotypes compared to the *tg1* and *tg2* (**Fig. 6-B**), reinforcing that the changes in the total protein content are due to the different concentration of SSPs across the explored genotypes. Curiously, it also indicates a similar trend for both lipids and pectin intensities.

**Figure 7** shows the synchrotron-based XRF mapping of the K, Zn, and Ca spatial distribution recorded in the same seed cotyledon tissue samples screened using FTIR. It shows that most of the Ca and K are found in the cell walls, whereas Zn is mainly located within the cotyledon cells (**Fig. 7-A**). Notable, clear Ca hotspots are also presented, which are likely associated with calcium oxalate crystals (Ilarslan *et al*., 2001).

**Figure 7.**
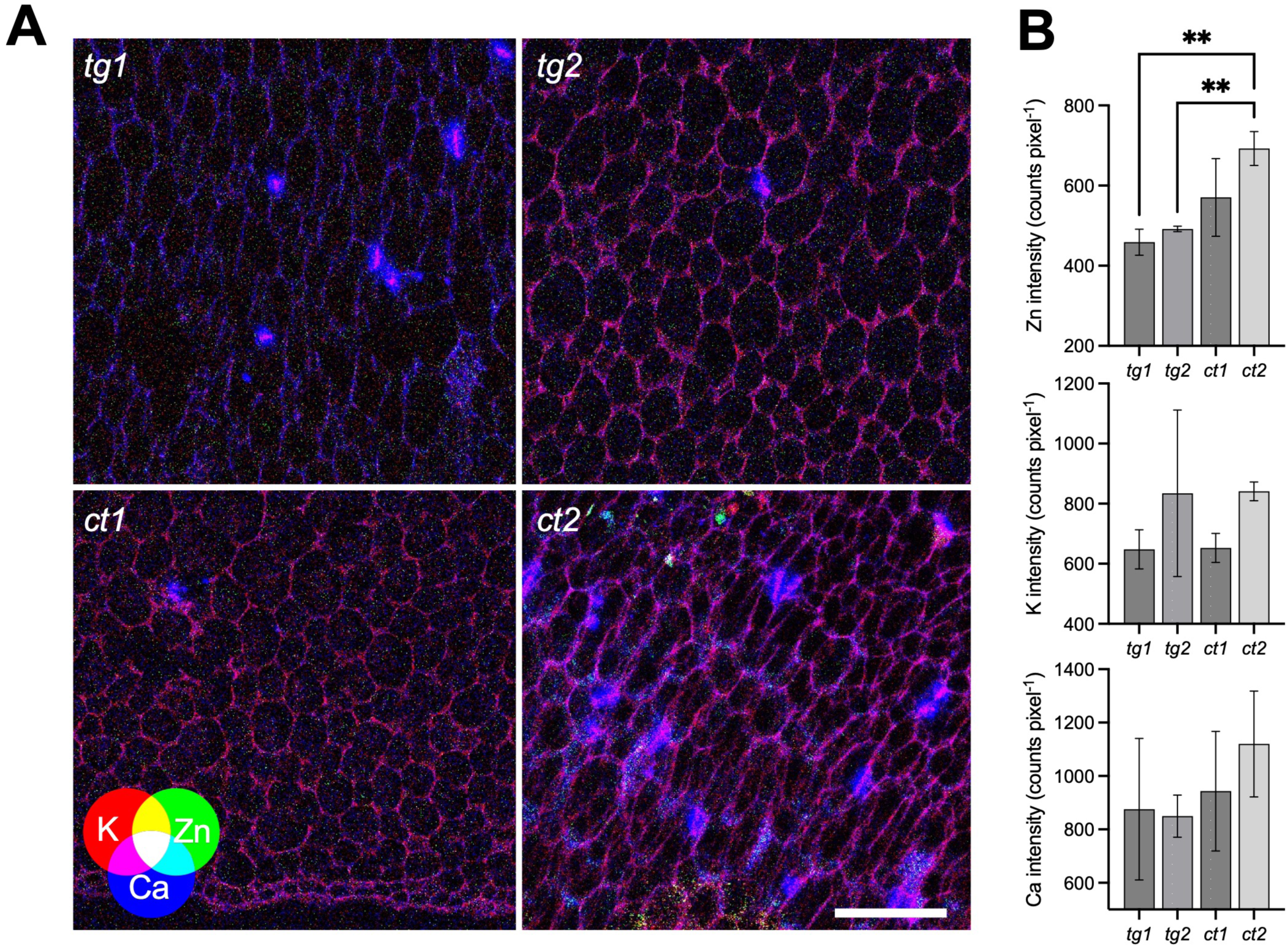
Spatial distribution (a) and average intensities (b) of K (red), Zn (green), and Ca (blue) XRF intensities recorded in the cotyledon tissue of mature (R_8_ stage) soybean seed cross-sections from varieties, *tg1*, *tg2*, *ct1*, and *ct2*, collected in field conditions. The bars represent the mean ± standard error recorded using at least 20 intracellular regions of three independent biological replicates. Data were subjected to Kruskal–Wallis one-way analysis of variance followed by Dunn’s multiple comparisons test at 0.05 level. ** = *p* <0.01. Scale:100 *μ*m

The subcellular quantification shows a higher Zn intensity in the *ct1* and *ctg2* seeds compared to the *tg1*, and *tg2* ones (**Fig. 7-B**), whereas the intensities of Ca and K did not significantly differ across the genotypes.

The characterisation of SSPs in soybean seed cotyledons was performed through light microscopy **(Fig. 8)**. As illustrated in **Fig. 8-A**, the 1% v v^-1^ xylidine ponceau staining shows that PSVs comprise most of the seed cotyledon cells. The quantitative analyses showed that *ca*. 40-50 PSVs of *ca*. 10 – 25 µm^2^ are found in each cell. Although the data suggest that the PSVs are larger in those genotypes presenting higher protein content, *i.e.*, *tg2* and *ct2*, the data exhibits a high variability, and no significant differences across the genotypes were observed (**Fig. 8-BC**). In addition, this quantification regards a two-dimensional perspective and does not provide any information regarding the PSV’s volume. This observation is related to the cotyledon cell morphology and the difficult to access SPPs volume with a micrometre section.

**Figure 8.**
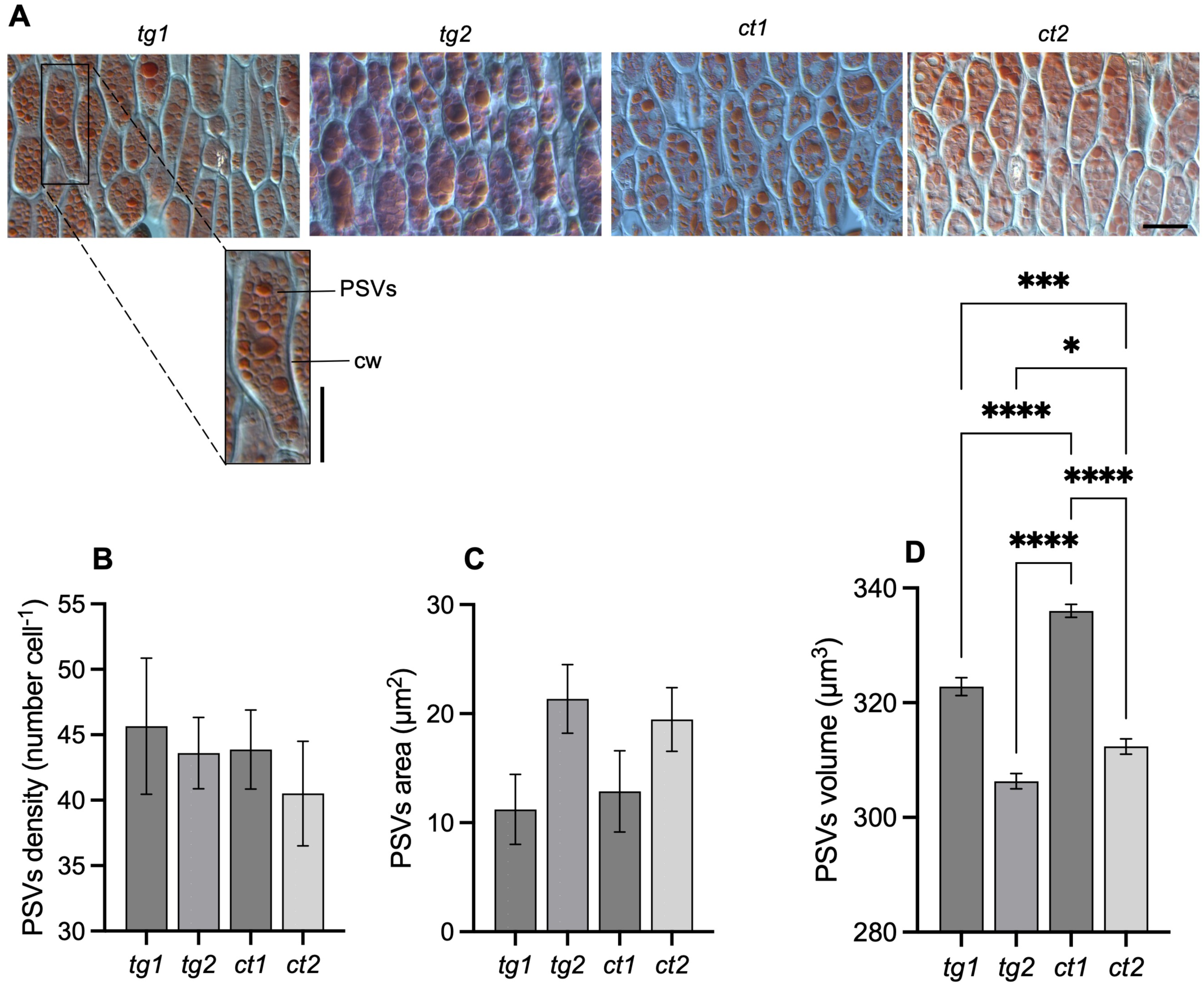
Light microscopy images detailing the protein storage vesicles (PSVs) in the cotyledonary cells of mature soybean seeds from *tg1*, *tg2*, *ct1*, and *ct2*, collected in field conditions collected under field conditions (a). Quantification of the protein storage vesicles density (b) and area (c). synchrotron-based *μ*-CT quantification of PSVs volume (d). Data represents the mean ± standard error recorded on at least 3 independent replicates and were subjected to Kruskal–Wallis one-way analysis of variance followed by Dunn’s multiple comparisons test at 0.05 level. The seed cross-sections were stained with 0.1% v/v xylidine ponceau solution for protein visualisation. cw: cell wall. Scale:100 *μ*m. *, **, ***, *****= *p* <0.05, *p* <0.01, *p* < 0.001, *p* < 0.0001.

In this scenario, we have evaluated the soybean seed cotyledonary tissues using a synchrotron-based computed tomography (**Fig. S2**), which enabled a proper segmentation and determination of the PSVs volume, as shown in **Fig. 8-D**. The µ-CT data reveals that *tg1* and *ct1* genotypes exhibited the largest PSVs volume compared to *tg2* and *ct2* ones, respectively. It means that the higher the protein content, the larger the protein bodies, suggesting that the number of PSVs is determined prior to the onset of the storage protein accumulation in seeds (Herman and Larkins, 1999).

These findings present a comprehensive characterisation of the dynamics of proteins and mineral nutrient accumulation, revealing that soybean seed genotypes exhibiting higher total protein content also present an increased abundance of SAA-rich (globulin) SSPs over the SAA-depleted one (β-conglycinin), as well as higher total S and Zn concentration. In addition, these seed genotypes also present increased protein and Zn content at the cellular level in the cotyledon cells, thereby reinforcing that Zn is associated with SSPs biosynthesis in soybean seeds and that Zn and S availability in the seeds might affect its protein content and profile. Future studies assessing whether the increase on both Zn and S availability changes SSPs content and profile are required to elucidate the potential extension of these approaches towards the enhancement of the content and quality of proteins in soybean seeds.

## 4. STATEMENTS

### Supplementary Data

Fig. S1. Details of the experimental plot where the four soybean seed varieties were sown.

Fig. S2. Workflow of synchrotron-based μ-CT data processing.

Fig. S3. Details of soybean cotyledon preparation for the sequential FTIR and synchrotron-based XRF mapping.

Fig. S4. FTIR spectra obtained in soybean seed cotyledon samples.

Fig. S5. Deconvoluted and fitted XRF spectra obtained in soybean seed cotyledon samples.

Fig. S6. Calcium and zinc quantification at the cell wall and cytoplasm of soybean seed cotyledons

Fig. S7. Accumulated macronutrients content throughout the development of the four soybean seed varieties used in the study.

Fig. S8. Accumulated maicronutrients content throughout the development of the four soybean seed varieties used in the study.

Fig. S9. SDS-Page gels from all the independent replicates.

Fig. S10. Normalised intensity of the 0-conglycinin and glycinin protein subunits from the four soybean seed varieties.

## Acknowledgements

This research used facilities of the Brazilian Synchrotron Light Laboratory (LNLS), part of the Brazilian Center for Research in Energy and Materials (CNPEM), a private non-profit organization under the supervision of the Brazilian Ministry for Science, Technology, and Innovations (MCTI). The Carnaúba and Imbuia beamlines staff, namely Dr. Raul de Oliveira Freitas and Dr. Ohanna Maria Medeiro da Costa, are acknowledged for their assistance during the experiments [proposals 20232796 and 20232798]. Dr. Renata Santos Rabelo is also acknowledged for the assistance with the sample preparation. Finally, we acknowledge the Biomedical Imaging and Therapy (BMIT-BM) beamline of the Canadian Light Source (CLS) in Saskatoon, Canada, where the µ-CT analyses were performed [proposal #39G13855]. An extend gratitude to our local contact, Dr. Jarvis Stobbs, for the support with the measurements and data processing. Dr. Fernando Guidorizzi and Linker Augusto Santos Louvison from ICL América do Sul are acknowledged for their support with the field trials. Dr. Marcos Rafael Petek from EMBRAPA Soja is acknowledged for providing us the conventional soybean varieties.

## Author Contributions

GSM and HWPC: conceptualization; GSM and HWPC: methodology; GSM, LCP, AL, CL, DM, ES, FSF, GPL, NGCS, JPRM, EA, AV, and CAP: formal analysis; GSM, LCP, AL, and ES: investigation; GSM, PV, SS, FSL, and HWPC: resources; GSM: data curation; GSM: writing - original draft; GSM, AL, CL, DM, JPRM, PV, and HWPC: writing - review & editing; GSM: visualisation; SS, FLS, and HWPC: supervision; GSM and HWPC: funding acquisition

## Data Availability

Raw data will be shared upon request to the corresponding author.

## Abbreviations

Cys: cysteine
2ER: endoplasmic reticulum
FTIR: Fourier transform infrared spectroscopy
ICP OES: inductively coupled plasma optical emission spectrometry
Met: methionine
PSVs: protein storage vacuoles
QTLs: quantitative trait loci
SAA: sulphur amino acids
SSP: seed storage proteins
XRF: X-ray fluorescence spectroscopy

